# Prosocial Decision-Making and Human Brain: A Graph Theory Analysis on Resting-State Functional Magnetic Resonance Image

**DOI:** 10.1101/2023.09.01.555856

**Authors:** Jinwei Xu, Delin Sun

## Abstract

Prosocial behavior is the cornerstone of a harmonious society. However, the functional organization of the brain underlying prosocial decision-making needs to be further explored. Here, we used graph theory analysis to investigate the brain functional connectivity derived from functional magnetic resonance imaging (fMRI) scans during rest in 55 female Chinese university students. The behavioral responses were collected in another fMRI scan during which participants completed an economic exchange game task by making multiple choices between a prosocial and a selfish option against either human counterparts (i.e., social environment) or robot counterparts (i.e., non-social environment). We found that making more prosocial decisions is accompanied by a longer path length in the right anterior superior temporal gyrus (aSTG), higher degree centrality in the posterior cingulate cortex (PCC), and higher betweenness centrality in the left aSTG. Our results suggest that human prosocial decisions are associated with greater inter-regional collaborations that are dominated by a few core nodes within the brain network of *Theory of Mind* (ToM). Moreover, an individual’s preference for making prosocial decisions could be uncovered by graph theory analysis of the functional brain network even without explicit task requirements.

## 1. Introduction

Value-based decision-making is common in nature. It occurs when we have to choose an option from several alternatives based on subjective values (i.e., purely considering our values and preferences), and can therefore also be called individual decisions (Rangel et al., 2018). However, the majority of our important decisions are made in the context of social interactions because of our very complicated social environment (Rilling et al., 2011). According to Fehr and Camerer (2007), social decision-making can be defined as those that affect both oneself and others, and are therefore typically determined by the preferences of the self and others. Prosocial decisions have more to with intentions and desire attributions than selfish actions (Sun et al., 2018). Selfish decisions are more likely to consider only self-interest, while prosocial decisions are more likely to consider counterparts’ interests along with their own (perhaps to get long-term and more benefits, or to seek stable relationships, etc.). There are many kinds of prosocial behaviors, including trust, reciprocity, altruism, fairness, social norm compliance, etc. (Rilling et al., 2011). Long-term stable cooperation requires altruism, which foregoes short-term rewards in exchange for a solid alliance that will provide long-term benefits (Fehr et al., 2007). Undoubtedly, prosocial decision-making is important to help people understand and anticipate social changes, prevent or reduce social problems, and ultimately promote peaceful social progress (Rilling et al., 2011; Rangel et al., 2018). A society without prosocial behavior will become unstable, and social problems such as violence will emerge. For example, tragic tragedies like car accidents will occur if people do not obey traffic rules. People won’t be able to reach a good cooperation relationship, which may stall their work process if they just consider their interests while ignoring the interests of others.

The term *Theory of Mind* (ToM), which describes the ability to infer the intentions of others, was coined by Baron-Cohen et al. in 1985. Understanding others’ needs and thoughts may enhance participation in prosocial activity (Dunfield et al., 2014), which may benefit others and help contribute to the development of a better ToM (Weller et al., 2014). Superior temporal gyrus (STG), posterior cingulate cortex (PCC) and medial prefrontal cortex (mPFC) are brain areas related to ToM (Sun et al., 2018), implying that these regions play significant roles in inferring and interpreting the intentions of others. The STG is involved in the perception of facial expressions (Sun et al., 2018), and has been discovered to be an important structure in social cognition processes (Bigler et al., 2007). The right STG plays an important role mainly in social perception and the left STG plays not only in social perception but also in language processing (Roger et al., 2010). The STG is associated with prosocial emotions in several studies, with the prosocial emotions of guilt, embarrassment, and empathy activating the bilateral STG and medial prefrontal cortex (mPFC) harmful consequences for others and for oneself as a result of one’s actions are associated with stronger feelings of guilt and strong activation of the right STG, posterior cingulate cortex (PCC) and medial prefrontal cortex (mPFC) (Moll et al., 2007; Morey et al., 2012). According to numerous reports, the right STG is involved in detecting, predicting, and reasoning about the motives and behaviours of others (Allison et al., 2000). Studies have showed that the right STG is more strongly activated while making prosocial decisions than when making selfish ones (Sun et al., 2018). In a study about the neural correlations in social cognition and empathy of borderline personality disorder, brain responses during cognitive empathy were significantly reduced in the left STG in patients with BPD (Dziobek et al., 2011). The PCC, on the other hand, is involved in self-referential processing (Schurz et al., 2014), for instance, in order to infer the intentions of others when perceived gaze shifts imply a lack of interpersonal engagement, people must think back on their personal experiences, which activates the PCC. For example, in Rilling et al.’s (2004) economic game task, stronger activation of PCC, mPFC, and temporoparietal junction (TPJ, which overlaps with STG to some degree) was detected when participants inferred the intentions of human counterparts through feedback. PCC is involved in a wide range of highly integrated tasks related to self-relevant processes, including self-expression and self-reflection, awareness, and other related processes. Also, the PCC appears to play a central role in networks related to social cognitive processing by generating excitatory effects on all other nodes (Esmenio et al., 2020). Therefore, studying the neural processing in the ToM-related STG, PCC, and mPFC when making social decisions may better explain how people infer and understand others’ intentions and thus engage in prosocial behavior.

Blood oxygen level-dependent (BOLD) contrast mechanisms are used in functional magnetic resonance imaging (fMRI), a form of imaging technique, to display regional and temporal changes in brain responses (Wang et al., 2018). fMRI is widely used in fields such as cognitive neurology because of its wide availability, non-invasiveness, relatively low cost, and good spatial resolution (Glover et al., 2011). Through a process known as the hemodynamic response, neurons are given more energy by nearby capillaries when they are stimulated, giving them more regional cerebral blood flow and a larger supply of oxygen than they typically require. Blood oxygen level-dependent (BOLD) contrast imaging is the term used to describe this imaging technique (Wang et al., 2018). Participants are expected to complete a cognitive task while undergoing fMRI scanning in the majority of task-related fMRI research. However, fMRI cannot be used to study the activity of the brain regions in particular demographic groups because they may not be able to perform certain tasks, such as infants, people with Alzheimer’s disease, and persons with other clinical problems (Canario et al., 2021). For this reason, resting-state fMRI (rs-fMRI) is a variant of task-based fMRI that maps brain function by looking at brain signals at rest. Rs-fMRI focuses primarily on measuring the spontaneous activity of BOLD signals, measured at rest, without participants performing explicit tasks that may alter brain activity (Wang et al., 2018). Compared to task-based fMRI, rs-fMRI has several advantages. It is simpler and, because it is performed at rest, it can detect global trends in the brain that are not found with task-based fMRI (Canario et al., 2021). It is more accessible to certain groups, such as babies or the elderly, because they do not need to perform movements that they may have difficulty imaging. Schizophrenia, bipolar illness, autism spectrum disorder, and attention deficit hyperactivity disorder are just a few of the mental disorders that have been extensively studied with this method (Canario et al., 2021). Furthermore, rs-fMRI has been found to pick up brain trends that task-based fMRI cannot or does not pick up (Wang, et al., 2018). Moreover, findings found by rs-fMRI have higher replicability compared to task-based fMRI (Viviano et al., 2018). One of the computational approaches to brain connectivity is functional connectivity (FC) (Canario, 2021). Functional connectivity studies the statistical dependence or temporal correlation between neurophysiological events that are spatially distant from each other (Canario, 2021). Graph theory analysis, seed-based functional connectivity analysis and other techniques can all be used to explore and identify functional connection (Wang et al., 2018). A functional connection (FC) considers the association between a given pair of brain regions of interest (ROI). All of the FCs across the whole brain region pairs form a brain network in which each ROI is a node and each FC is an edge. Graph theory further simplifies the brain network by turning all suprathreshold edges into 1 (connected) and the rest edges into 0 (unconnected), and is widely used to investigate the characteristics of the whole brain network as well as the roles of each node in this network (Shinn et al.,2021). Several preprocessing steps are performed on the acquired fMRI data, averaging the time course of all voxels within a selected specific region as a time series for that brain region. Next, pairwise correlations (correlation matrices) of the time series between different brain regions of the brain are determined using methods such as correlation analysis. Once the correlation matrix has been supra-threshold connected depending on connection strength or network sparsity, the binary connection matrix (also known as the adjacency matrix) is obtained. Finally, the CONN functional connectivity toolbox can be used to extract essential topological features that characterise the local and overall architecture of brain network connectivity (**Fig.1**) (Shinn et al., 2021). In graph theory, various parameters can be used in the overall analysis. These parameters include path length, degree centrality, clustering coefficient, betweenness centrality, global efficiency, and local efficiency (Canario, 2021; Wang et al.,2018). Of all the parameters of graph theory, the path length is the one we are most interested in. The path length is the number of edges connecting two nodes, a shorter path length between two nodes indicates a faster and more automatic transfer of information between the two nodes (Farahani et al., 2019). On the contrary, a longer path length indicates that the information transmission needs to travel longer and is slower (due to the collaborative processing across more brain nodes).

**FIGURE 1.**
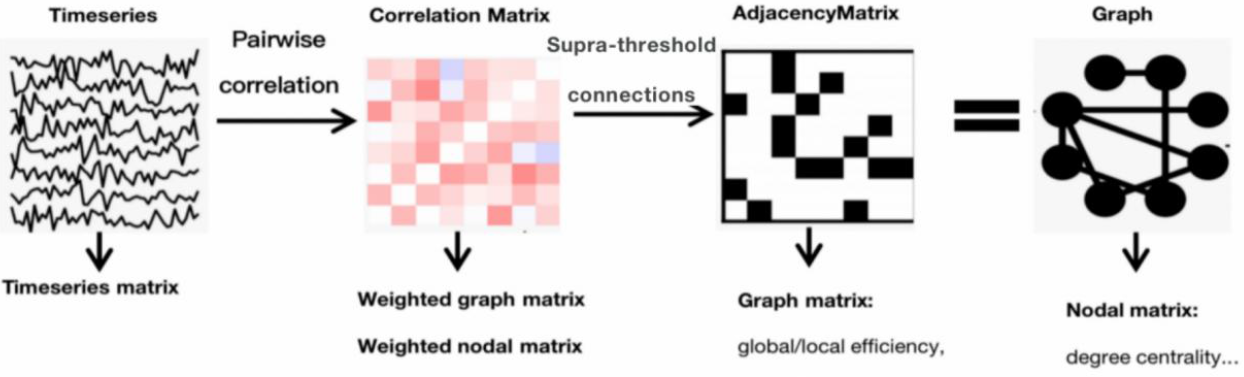
Diagram describing how to convert from fMRI regional time series to binary brain graphs. Pairwise correlations of time series between different brain regions of the brain (correlation matrices) were determined using methods such as correlation analysis. Then, the values of the correlation matrix are thresholded and weighted to filter out weakly connected networks and leave only strongly connected networks as a binary brain map (binary connectivity matrix, i.e., adjacency matrix). Finally, key topological properties describing the local and overall structure of the brain network connections (e.g. degree centrality, etc.) can be obtained using the CONN functional connectivity toolbox.

There are many studies investigating the brain functional connectivity of prosocial decision-making by using rs-fMRI. The vmPFC, temporal poles, and PCC, which are brain regions associated with self-referential thinking and reward processing, have stronger connectivity between them in the resting state, which makes individuals prone to honest behavior (Speer et al., 2022). FCs in social brain networks, which include brain regions like the PCC and the orbitofrontal cortex, are particularly predictive of dishonesty rates, as are FCs within reward networks and across self-referential and cognitive control networks (Pang et al., 2022). These studies have explored the functional connectivity of brain regions involved in individual decision-making during resting states, providing additional evidence to explore social decision-making. However, their studies are limited in that they focus on the activity of specific relevant brain regions and do not explore the overall functional connectivity of the whole brain during prosocial behavior. Most studies have been more concerned with the strength of the functional connectivity or activations of the brain regions or networks involved in prosocial behavior, but few have explored the path lengths of the functional connectivity embodied in these brain regions, i.e., the efficiency of the speed of information transfer. We are curious whether ToM-related brain regions such as STG and PCC have higher information transfer efficiency when making prosocial decisions and can be used to predict more prosocial behaviors. Using path length might determine whether the transfer of information between the STG and other relevant brain regions of the ToM is more efficient and faster when making prosocial decisions. In this study, Participants underwent task-related and resting-state fMRI scans. During a novel economic exchange game task, they made selfish or prosocial decisions when against a robot counterpart (i.e., non-social environment) and against a human counterpart (this situation is elicited by displaying a photo of a pair of human eyes on the screen) (Sun et al., 2018). The resting-state scans were performed after the task-relevant scans were completed. Therefore, we propose two alternative hypotheses: (1) Human brain is wired for selfishness. Therefore, more selfish people have shorter connections between the STG and other brain regions. (2) Human brain is wired for prosocial actions. Therefore, more prosocial (less selfish) people have shorter connections between the STG and other brain regions.

## 2. Methodology

### 2.1 Participants

East China Normal University recruited fifty-five healthy female university students to take part in a study looking at the brain correlates of social decision-making. The study was carried out by the Declaration of Helsinki’s ethical principles, and each participant supplied written informed permission. The details of this study have been previously published (Sun et al., 2015; Sun et al., 2016; Sun et al., 2018), and the present study is a secondary analysis of the existing data without any collection of new data from the same participants or recruitment of new participants.

### 2.2 Procedures

As illustrated in **Fig. 2**, participants saw pictures of human eyes (direct or averted gaze, which was not differentiated in the present study) and robot’s eyes in random order during a task-related fMRI scanning before the resting-state fMRI scanning. And the positions of the bars indicating the two alternatives (left and right) were balanced. On each trial, participants received the counterparts’ financial contribution along with an offer for how to divide the remaining funds (the allocation amount was a randomly generated number between 80 and 150, and the suggested return to the counterparts was a percentage of 60%, 65%, or 70%). Every trial requires the participant to make a decision by pressing one of two buttons with their right index or middle finger for 3 seconds (the decision phase) during which time the increased investment amount is displayed on the screen. All human players will finally get a real money bonus based on how much they made while completing the task. The participants have the option to accept or reject the offer, selecting a different course of action that is better for them but worse for their rivals. The counterparts will then have a 50% chance of determining whether the participant accepts or rejects the offer. If the participant reject the offer and be successfully detected by the counterpart, the portion of rewards the participant in the trial will be shifted to her counterpart. The participant will keep the share if the refusal is not noticed. If the participant does not act, the counterpart in that trial will receive all benefits.

**FIGURE 2.**
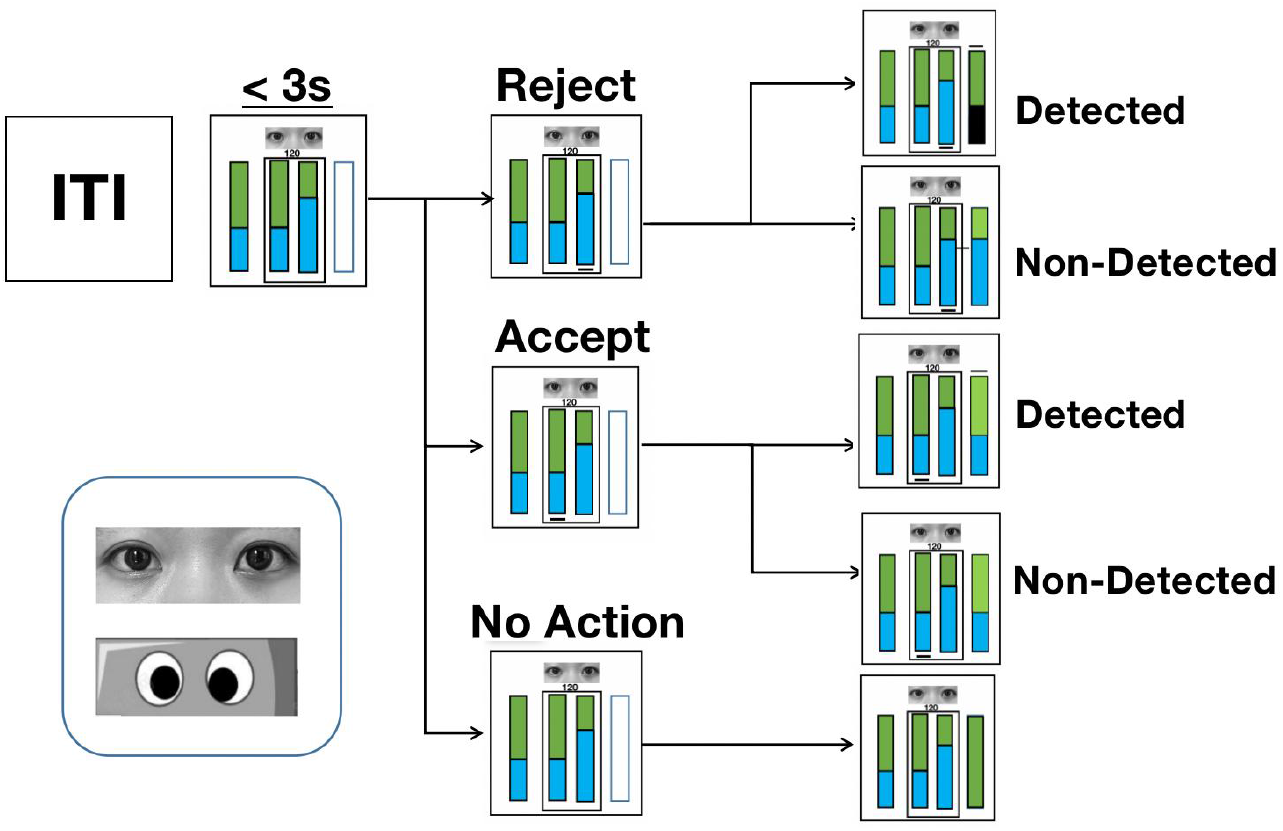
Task paradigm and eye stimuli. The proportion of the reward given to the counterpart and the participant is indicated by the green and blue parts of the vertically stacked bars, respectively. In each trial, after learning the total amount to be divided (i.e., the number on the screen) and the counterpart’s offer (represented by the bar on the left), the participant could accept or reject the offer by pressing one of the two buttons corresponding to the bar in the middle of the screen within 3 seconds. Accepting the counterpart’s offer is beneficial to both of them, however, rejecting the offer indicates a strategy that is more favorable to the participant. The associated bar graph displays a black line immediately after the selected action. The results bar on the right side shows the final reward distribution for the trial in the last three seconds. If the participant’s choice is successfully detected by the counterpart, a black line is shown above the result bar. Conversely, if the detection fails, no line is displayed. When a refusal was detected, the participant will lose her portion of rewards on the trial and her portion of the result bar turned black. In the other conditions, the participant can keep her portion. If there was no response or the response took longer than the three-second decision period, the reward for the trial was given to the counterpart. The average time between trials (ITI) and between stimuli (ISI) was three seconds. At the beginning of each trial, one of two cues was presented. The robot’s eyes and the human’s eyes were the cues.

### 2.3 Data Acquisition

Using a 3-Tesla Siemens Trio MR scanner, brain imaging data, including structural images, task-related functional images, and resting-state functional images, were gathered. 35 axial slices covering the entire brain were used to create functional pictures using a T2*-weighted echo planar imaging (EPI) procedure. Using 3D MRI sequences, a high-resolution structural picture of each individual was also obtained. In this order, participants underwent structural scanning, resting-state scanning, and task-related scanning. Participants were told to keep their eyes open during the resting-state scanning for roughly 10 minutes without any specific activity requirements.

A task-related scanning procedure was used to gather behavioral data utilizing an integrated functional imaging system (IFIS), which allowed for the display of visual stimuli as well as the recording of responses. The details of the decision-making task have been reported in previous studies (Sun et al., 2015; Sun et al., 2016; Sun et al., 2018), and in summary, participants played an economic decision-making task in which they were presented with a choice between a safe choice that mildly increased monetary reward without risk and a risky choice that had the potential to largely increase the reward but also included a large loss. During trials in which participants played against a computer, they only had to consider their benefits, which was defined as individual decision-making. In trials against human counterparts, they were asked to consider both their interests and those of other human players, which was defined as social decision-making. In both the individual and social decision-making contexts, the frequency of making a risky or safe choice as well as the reaction time (RT) was measured.

### 2.4 Image Data Preprocessing

The CONN functional connectivity toolbox, implemented in MATLAB, was used to preprocess image data. Segmentation, normalization to Montreal Neurological Institute (MNI) space, resampling at 2-mm3, motion and slice timing correction, functional outlier evaluation, and smoothing with a Gaussian kernel were all steps in the preprocessing procedure. Denoising was used to scrub variables, BOLD signals, and white matter and cerebrospinal fluid masks to eliminate putative realignment confounders. Additionally, utilizing band-pass filtering between 0.008 Hz and 0.09 Hz, linear trends were removed, low-frequency drift was abolished, and high-frequency respiratory and cardiac noise were both reduced.

### 2.5 Graph Theory Measures

#### Path Length

It describes the number of edges that must be traversed to get from one node to another. That is the path length that needs to be taken from one node to another. The shorter the path length that needs to be traversed, the faster and more efficiently the node pair can transfer information between them (**Fig. 3A)**.

**FIGURE 3.**
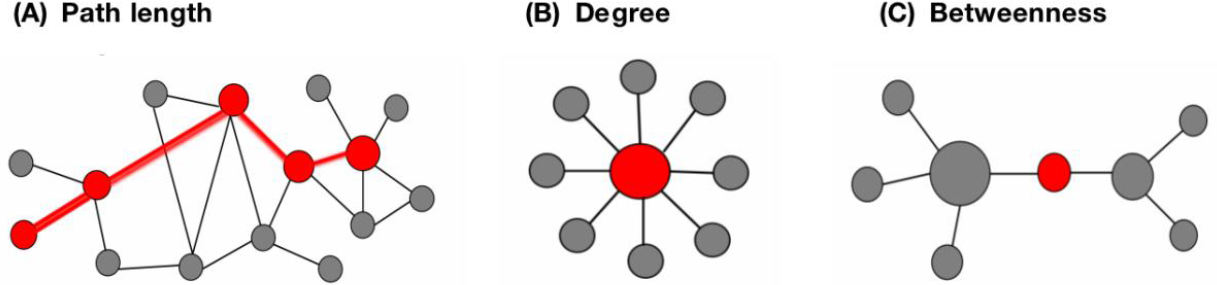
Brain Graph Measures Employed in This Study. **(A)** The path length measures the number of edges that must be traversed to get from one node to another. **(B)** The degree centrality is defined as the number of the node’s neighbors. **(C)** The betweenness centrality measures the node’s role in acting as a bridge between separate clusters by computing the ratio of all shortest paths in the network that contain a given node.

#### Degree Centrality

It is defined as the number of links incident upon a node (i.e., the number of edges that a node has). The degree can be interpreted as the direct risk of a node capturing any information flowing through the network. A node with a higher degree centrality may play a more important role in integrating information from diverse sources in the network (**Fig. 3B**).

#### Betweenness Centrality

It is a centrality metric in a graph based on the shortest pathways (**Fig. 3C**). There exists at least one shortest path between each pair of nodes in a connected graph that minimizes the sum of the edge weights or the number of edges the path passes over. The betweenness centrality of a node is determined by the quantity of these shortest paths that pass through it. The degree to which nodes are separated from one another is represented by betweenness centrality. As more information will move via that node based on the shortest path (therefore the most direct and fastest), a node with a higher betweenness centrality may have more control over the network.

### 2.6 Statistical Analyses

In this study, we aimed to investigate the neural correlates of two behavioral performances: (1) the frequency of making a selfish choice (Fre_Selfish), which is the contrast between the frequency of making a risky choice in social decision-making and the frequency of making a risky choice in individual decision-making; (2) the RT of making a selfish choice (RT_Selfish), which is the contrast between the RT of making a risky choice in social decision-making and the RT of making a risky choice in individual decision-making. The behavioral data obtained in the study have been elaborated in previously published papers. For selection frequency, no significant results were found for counterparts, selection, and their interaction. On the other hand, for reaction times, we found longer reaction times for human counterparts than for computer counterparts. No significant results were found for the effect of choice or the interaction between counterpart and choice (Sun et al., 2015; Sun et al., 2016; Sun et al., 2018). Therefore, in this study, we only analyzed fMRI results. With age included as a covariate of no relevance, we investigated the associations between behavioral performances and predefined neural measures using a linear regression model set up using the CONN toolbox. All statistical results were considered significant at the alpha level of .05 (two-tailed). Due to our strong research ideas regarding the included neural measures and the relevant brain nodes, we revised the number of sides (i.e., 2 for the left and right hemispheres) but not the number of all potential neural measures or the number of brain nodes. This method also avoids type II errors and facilitates future studies based on our pilot findings.

## 3. Results

### 3.1 Rs-fMRI Findings

The results of rs-fMRI analyses were shown in **Fig. 4**.

**FIGURE 4.**
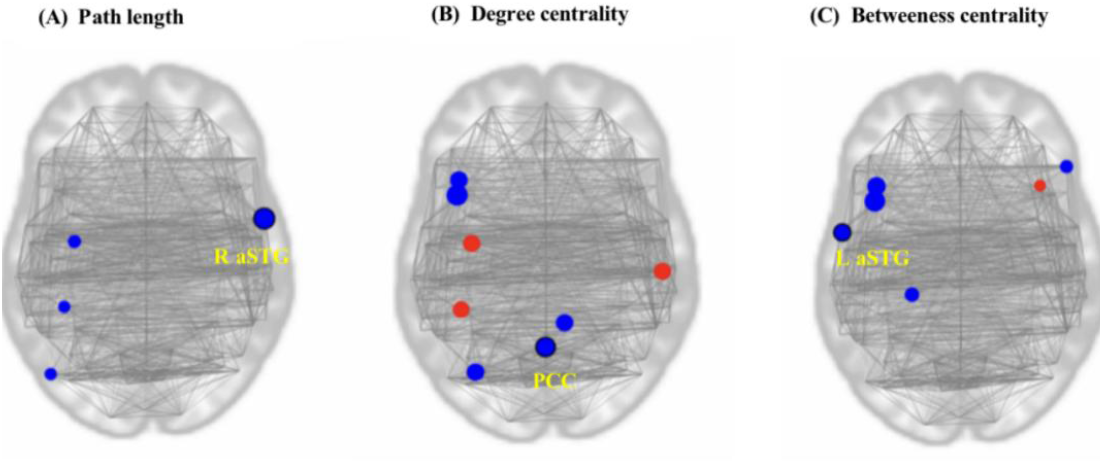
The resting-state fMRI results of selfish decision-making in this study. The red dots represent positive values, indicating a positive relationship between the graph theory measure in this region and the frequency of making selfish decisions (a negative correlation with the frequency of prosocial decisions). Blue dots represent negative values, indicating a negative relationship between the graph theory measure in this region and the frequency of making selfish decisions (a positive correlation with the frequency of prosocial decisions). **(A)** A higher frequency of making prosocial decisions is accompanied by longer path length in the right aSTG. **(B)** A higher frequency of making the prosocial decisions is associated with a higher degree centrallity in PCC. **(C)** A higher frequency of making prosocial decisions is associated with higher betweenness centrality in left aSTG.

#### Path Length

A longer path length in the right aSTG (T value = -3.40) is accompanied by a higher frequency of making prosocial decisions (**Fig. 4A**).

#### DegreeCentrality

A higher frequency of making prosocial decisions is accompanied by higher degree centrality in PCC (T value = -2.34) (**Fig. 4B**).

#### Betweenness Centrality

A higher betweenness centrality in the left aSTG (T value = -2.97) is accompanied by a higher frequency of making prosocial decisions (**Fig. 4C**).

## 4.Discussion and Conclusion

### 4.1 Discussion

Our study used resting-state fMRI to investigate the associations between brain graph measures and participants’ prosocial decisions. We found that the right anterior superior temporal gyrus (aSTG) showed longer path length with other brain regions when the frequency of making prosocial decisions is higher. We also found that a higher degree centrality in the posterior cingulate cortex (PCC) is associated with a higher frequency of making prosocial decisions. Moreover, higher betweenness centrality in the left anterior superior temporal gyrus (aSTG) is associated with a higher frequency of making prosocial decisions.

In our study, we found that the longer path length in the right aSTG is associated with a higher frequency of making prosocial decisions. It indicates that the right aSTG has a longer pathway to transfer information with other brain regions when making prosocial decisions, which means that cognitive resources located in more diverse brain regions may be needed to jointly make a more sophisticated choice regarding the interests of both self and others. This result is consistent with our first prior hypothesis, which is that the human brain is designed for selfishness and that more selfish people have shorter connections between the right STG and other brain regions. According to numerous studies, the right STG plays an important role in detecting, predicting, and reasoning about the motives and acts of others (Allison et al., 2000). And a stronger activation of the right STG was found in prosocial choices than in selfish choices (Sun et al., 2018). When making prosocial decisions, the right STG is more involved in making inferences about the other’s intentions, while the opposite is true when making selfish decisions. However, the current graph measures are based on whole-brain connections (Farahani et al., 2019), and our findings can only show that the right STG has longer pathways to other regions of the brain when making prosocial decisions, and it is not possible to know whether more connections exist with ToM-related regions such as the TPJ. Future work is needed to show whether there are more connections between ToM-related regions. Compared to choices related to self-interest only, making prosocial decisions require further detection and reasoning of the counterpart’s intention, and thus involves more brain regions in the collaborations with the right STG. The information flow may pass through more brain nodes to finish more complicated and sophisticated calculations. Therefore, it is reasonable to find that more prosocial actions are accompanied by a longer path length of information transfer.

The PCC has been reported to play roles in self-referential processing, and in encoding the neural processing of attention (Sun et al., 2015). Furthermore, we found that a higher degree centrality in PCC is associated with a higher frequency of making prosocial decisions. The results suggest that the PCC plays a more important role in integrating information from relevant brain regions when making prosocial decisions. When making prosocial decisions, there may be more processing between oneself and others. This finding is consistent with earlier research that people need to recall their personal experiences in order to infer the intentions of others while making prosocial decisions (Sun et al., 2018) since PCC is associated with episodic memory (Sun et al., 2015c). The more like the other person is to oneself, the more likely we are to draw inferences about them based on what we know about ourselves, claim simulation theories of social cognition (Esmenio et al., 2020). And Esmenio et al. claimed that PCC appears to play a central role in networks related to social cognitive processing by generating excitatory effects on all other nodes. Our findings are consistent with previous findings that when people make prosocial decisions, we mobilize more of our own experience to detect more of each other’s intentions because the other person is similar to us, and PCC needs to pool more information from relevant brain regions to engage in social cognitive processing. According to Mohanty et al. (2008), the PCC showed stronger activation in response to a rewarding stimulus compared to a stimulus without rewards. Because of the need to consider the interests of others more, the PCC shows stronger activation in response to reward stimulation, and at the same time awakens relevant brain regions to recall personal experiences and quickly integrates information transmitted from these brain regions to infer the intentions of the others, thus showing higher degree centrality in making prosocial decisions. More prosocial decisions can be made if more personal experiences are available.

Several previous studies showed that bilateral STG is associated with mentalizing and intention processing in social brain networks (Chen et al., 2022). In 2011, Dziobek et al. made the claim that patients with BPD had significantly reduced left STG brain responses during cognitive empathy. The left STG may be more involved in the cognitive processing of empathy because patients with BDP have deficits in both cognitive and emotional empathy. In our study, we found that a higher betweenness centrality in left STG is associated with a higher frequency of making prosocial decisions, which means the left STG may have more control over the relevant network of prosocial decision-making because more information will pass through it based on the shortest path (therefore the most direct and quickest). Our study shows that when making prosocial decisions, the left STG is one of the many information transmission pathways in the relevant network that must be passed through, and that many of the fastest and most direct pathways in the network connections of brain areas related to prosocial decisions must involve empathy, i.e., the more empathetic one can be with the other person, the more likely one is to engage in more prosocial behavior. While ToM and empathy-related neural networks are carried out by separate, independent brain networks, they are co-activated and engage in social interaction in a complex setting (Preckel et al., 2018). Our findings are consistent with the above view that the left STG region associated with empathy becomes a necessary route for information transfer from many relevant networks when making prosocial decisions. That is, because prosocial decisions are made with more consideration of the other person’s interests than one’s own, greater empathy is associated with a cognitive understanding of others’ thoughts and intentions.

The human brain is wired for selfishness. More selfish people have shorter connection paths in the right STG to other brain areas, meaning faster responses to make selfish decisions. Therefore, more effort is required from relevant brain regions when making prosocial behaviors. In making prosocial decisions that require more detection and reasoning about each other’s intentions, the right STG has a longer connection path to other brain regions. In making prosocial decisions, the PCC is more efficient at pooling information from peripherally relevant brain regions because people considers personal interests more than those of others, and more attention and speculation about the other’s interests are accompanied by more needed self-reference. Finally, since the left STG is involved in empathy-related neural processing and is accompanied by more empathy when considering the interests of the other person, the left STG becomes an important node in the information transfer pathway of the relevant brain regions at this time when more prosocial decisions are made. Therefore, its importance is stronger during prosocial decision-making. The PCC showed more connections to peripheral brain regions and the left STG played a mediating role in more node pairs, both results implying that more brain regions were involved when making prosocial decisions. And the longer pathways between people’s right STG and other brain regions under prosocial behavior (longer pathways indicate more brain regions involved) are consistent with the first two results. That is, more prosocial decision-making may require more relevant brain regions to be involved in collaboration.

Furthermore, our findings have clinical implications, particularly for those with autism spectrum disorders (ASDs) who have difficulty with reciprocal interactions with others (Baron-Cohen et al., 1997). Because patients with ASD have impairments in reciprocal cognitive processes, their relevant brain network connections cannot be explored by doing the task. Our study provides a safer and more accessible neuroimaging method to study individual preferences for prosocial behavior in people with ASD.

## 4.2 Limitations

Our study also has several limitations. First, our participants are all females, so the unbalanced gender distribution may hinder the generalization of the results. Future research can directly examine the neural correlates of males and females to completely understand how the brain processes prosocial decision-making. Secondly, our participants were all Chinese. People from different cultures may have different processing in prosocial decision-making. Future studies are needed for comparing brain functional connectivity across cultures. Third, only a small range of age participants are considered in our study. However, neural processing of social decision-making may differ by age. Since personal experience is one of the important factors influencing decision-making, the richer life experience of older people may lead them to be more inclined to make rational and beneficial choices in making social decisions than younger people (Zakirov et al., 2020). It’s interesting to compare the neural processing of prosocial decision-making across ages.

## 4.3 Conclusions

In our study, we explored the functional connectivity of brain regions associated with ToM in resting state, particularly the bilateral STG and PCC in making prosocial decisions. We found that the higher frequency of making prosocial decisions is associated with a longer path length in the right STG, a higher degree centrality in PCC, and a higher betweenness centrality in the left STG. Our findings suggest that people are quicker and more automatic at making selfish decisions, while stronger functional connections and engagement of ToM-related brain regions are required for making prosocial decisions. People with a stronger preference for making prosocial decisions are accompanied by greater inter-regional collaborations in the brain network, particularly significant at brain nodes that play important roles in ToM. Furthermore, our study suggests a potential clinical application in inferring an individual’s preference for prosocial actions based on a safe and easy-acquired neuroimaging method.

